# Single-cell transcriptome analysis reveals the cellular atlas of human intracranial aneurysm and highlights inflammation features associated with aneurysm rupture

**DOI:** 10.1101/2023.04.06.535955

**Authors:** Hang Ji, Yue Li, Haogeng Sun, Ruiqi Chen, Ran Zhou, Anqi Xiao, Yongbo Yang, Rong Wang, Chao You, Yi Liu

## Abstract

Intracranial aneurysm (IA) is pouch-like pathological dilations of cerebral arteries, which often affects middle-aged people and culminates in life-threatening hemorrhagic stroke. A deeper knowledge of the cellular and gene expression perturbations in human IA tissue deepens our understanding of disease mechanisms and facilitates developing pharmacological targets for unruptured IA. In this study, 21,332 qualified cells were obtained from cell-sparse ruptured and unruptured human IA tissues and a detailed cellular profile was determined, including conventional endothelial cells, smooth muscle cells (SMC), fibroblasts and the newly identified pericytes. Notably, striking proportion of immune cells were identified in IA tissue, with the number of monocyte/macrophages and neutrophils being remarkably higher in ruptured IA. By leveraging external datasets and machine learning algorithms, a subset of macrophages characterized by high expression of CCL3 and CXCL3, and transcriptional activation of NF-κB and HIVEP2 was identified as the cell most associated with IA rupture. Further, the interactome of CCL3/CXCL3 macrophages disclosed their role in regulating vascular cell survival and orchestrating inflammation. In summary, this study illustrated the profile and interactions of vascular and immune cells in human IA tissue and the opportunities for targeting local chronic inflammation.

## Introduction

Intracranial aneurysms (IA) are common intracranial vascular lesions, often manifesting as confined pathological outpouchings of the Circle of Willis, with an incidence of approximately 3∼5% in the general population^1, 2^. Spontaneous subarachnoid hemorrhage (SAH) caused by IA rupture is extremely fatal and disabling, often affecting middle-aged people^3^. Worryingly, an increasing number of people are found to carry single or multiple unruptured IAs during routine screening with the spread of angiography^4^. Despite recent advances in the clinical management of IA, endovascular and surgical aneurysm repair, or simply conservative management have inherent risks of complications, making effective pharmacological intervention an appealing choice.

The development of promising pharmacological targets is based on a thorough understanding of the mechanisms of IA progression and rupture. Although the tiny, cell-sparse tissues derived from patients make research difficult, advances in technology have yielded important insights into the pathophysiological alterations of IA. Histological studies based on patients’ and murine model IA tissues highlight decellularization, internal elastic lamina remodeling, and chronic inflammation during the dilation of vascular wall^4, 5^. Magnetic resonance imaging-based studies suggest an association between circumferential aneurysm wall enhancement with inflammation and macrophage infiltration, which serves as a sign of increased risk of rupture^4^. Over the last decade, 3-dimensional (3D)- or 4-dimensional (4D)-flow-based studies have elucidated the hemodynamic perturbations associated with the onset and progression of IA, adding critical evidence for the morphological and functional conversions of endothelial and SMC induced by wall shear stress^5^. At almost the same period, genome-wide association studies based on large scale cohorts illuminate genetic risk locoes, genes, and modifiable epigenetic factors associated with IA pathology^6, 7^. Overall, these scientific advances have delineated an outline of the pathological mechanisms of IA.

Cells are the main building blocks of cerebrovascular system and their status has a profound impact on physiological function and disease progression. Recent studies based on single-cell transcriptome sequencing have established a comprehensive atlas of cerebrovascular cells and provided unprecedently detailed knowledge on cellular and gene expression in disease states^8, 9^. Further, a single-cell RNAseq-based study utilizing murine IA tissue pinpoints the role of macrophages in disease progression, which are also responsible for bleeding in intracranial arteriovenous malformations and thoracic aortic aneurysms^8,10, 11^. Cellular vulnerabilities of general human IA tissue and gene expression dysregulations associated with IA rupture are hitherto largely unknown. In this perspective, we performed single-cell RNA sequence and comprehensive bioinformatic analysis on patient-derived IA samples at different pathological stages to address fundamental molecular and cellular dynamics and inflammation features implicated in IA rupture. Our study provides a detailed reference of cell types and gene expression dynamics of human IA tissue and preliminary basis for the targetable components in local chronic inflammation.

## Materials and methods

### Enrollment of participants and collection of samples

The protocol of human IA tissue collection was permitted by the Ethics Review Committee of West China Hospital, Sichuan University. All participants were informed and signed a written consent form priorly. Experiments using human tissue were performed in accordance with the relevant guidelines and regulations. Patients were diagnosed as IA angiographically before bypass surgery or clipping. Fresh IA tissues were obtained from 3 SAH and 1 unruptured IA patients. Tissues were obtained in close collaboration with two senior neurosurgeons with extensive experience in IA surgery to minimize destruction. Thrombus-containing IA tissue was avoided. The obtained IA tissue was immediately flushed of erythrocytes. Patients were excluded if they had a heritable arterial disease (Marfan syndrome, Ebler-Danlos syndrome, pseudoxanthoma elasticum, autosomal dominant polycystic kidney disease, neurofibromatosis type I). Two external microarray-based IA datasets were retrieved from the Gene Expression Omnibus, including GSE13353 and GSE122897^12, 13^. Positive and negative hits of IL-4- and IFNγ-polarized macrophage, and gene signatures of different conditions-polarized macrophages were retrieved from GSE47189^14^.

### Single-cell RNA sequencing sample preparation

Fresh specimens were placed in chilled MACS tissue storage solution (Miltenyi, Germany) and transported to the laboratory on ice. The IA tissue was slightly flushed with prechilled DMEM (Sigma-Aldrich, USA) to remove erythrocytes. Then, the tissue was fragmented and digested in mixed digestion buffer (normal Dulbecco’s modified Eagle’s medium containing 15 mg/ml collagenase type II + 500 ug/ml elastase) for approximately 30 mins at room temperature. Tissue suspension was filtered through a sterilized a 70-μm followed by a 40-μm cell strainer to isolate debris. Isolated cells were collected by centrifugation at 500g for 5 minutes at 4℃, and the supernatant was carefully aspirated. The cell pellet was resuspended in Gibco ACK Lysis Buffer (Thermo Fisher Scientific, USA) for 3 minutes to lyse residual erythrocytes. Cells were then collected by centrifugation at 500g for 5 minutes and washed three times with sterile RNase-free phosphate-buffered saline (PBS) (Beyotime, China) containing 0.04% BSA (Beyotime, China). Thereafter, cells were pelleted and resuspended in PBS containing BSA. Cell viability and concentration were assessed automatically using the Countstar Rigel S2 system (Countstar China). The microfluidics-based cDNA library construction and quality control, and sequencing based on the Novaseq6000 platform were conducted by Biomarker Technologies CO., Ltd. (Beijing, China).

### Single-cell RNA sequencing data analysis

Count matrices of cell-by-gene was created using Cellranger (v7.1.0). Raw sequencing reads were aligned to the pre-mRNA annotated by homo sapiens reference genome GRCh38. Seurat framework was employed to manage the downstream analysis. DoubletFinder was employed for potential doublet detection at a criteria of 7.5% doublet formation rate based on the recommendation of 10× Genomics^15^. A minimum of 500 genes and 15% mitochondrial cutoff were used to remove low quality cells. The SCTransform workflow was used for count normalization. Diagonalized canonical correlation analysis (CCA) and mutual nearest neighbors (MNN) were performed for batch correction^16^. The scaled data with variable genes identified by Seurat were used to perform principal component analysis (PCA). The first 30 principal components of the PCA were used for clustering and cell identification. Pseudotime analysis was performed using the R package ‘monocle3’^17^. CellChat and cellphonedb (V3.0) were employed to predict cell-cell communication^18, 19^. SCENIC was used to identify activated transcription factors^20^. Weighted gene co-expression network analysis (WGCNA) was performed to identify the intrinsic gene expression program^21^. As the single-cell count matrix is sparse, pseudocells was constructed as the averages of 5-10 randomly chosen cell within each cluster. Functional enrichment analysis was performed using the webtool Metascape^22^ (https://www.metascape.org/).

### Correlation between immune cells with IA rupture

Marker genes of each type of cell were defined as those average log2FC > 0.5, Pct.2 < 0.15, and adjusted p-values < 0.1. Single-sample gene set enrichment analysis (ssGSEA) were employed to estimate the abundance of each type of cell in bulk sample. Receiver operating characteristic curves (ROC) and corresponding area under the curves (AUC) were calculated for estimating the correlation.

### Statistics

All statistics were performed using the R software (v4.2.1). Wilcoxon-test was employed for the comparison of non-normally distributed variables.

## Results

### Cellular and molecular profiles of IA

Raw data of the 4 samples was individually analyzed and embedded^16^. A total of 21,332 qualified cells were obtained for further analysis (Supplementary Figure 1A-C). Unbiased cell clustering analysis was performed and 21 clusters were initially obtained (Supplementary Figure 1D-F). Based on established cell identity specific transcriptomic biomarkers, 8 major cell types were first assigned (Figure 1A, B). Vascular cells included endothelial cell (CLDN5, PECAM1)^9^, pericyte (ABCC9, KCNJ8)^8^, fibroblast (DCN)^8^ and SMC (MYH11, CNN1)^11^, accounting for approximately 4.8% of the total cells. Immune cells made up the majority of the IA tissue, including granulocyte, mostly neutrophil (S100A8, CXCL8)^10, 23^, monocyte/macrophage/dendritic cell (CD14, CD16)^24^, T cell (CD3E, CD3G)^9^, NK (KLRF1, GZMA)^9^, B cell (CD79A, MS4A1)^9^ and mast cell (TPSAB1, CPA3)^9^. No type of cell was absent from a sample, while their proportions varied considerably in rupture (IA1∼3) and unruptured (IA4) tissues (Figure 1C).

**Figure 1.**
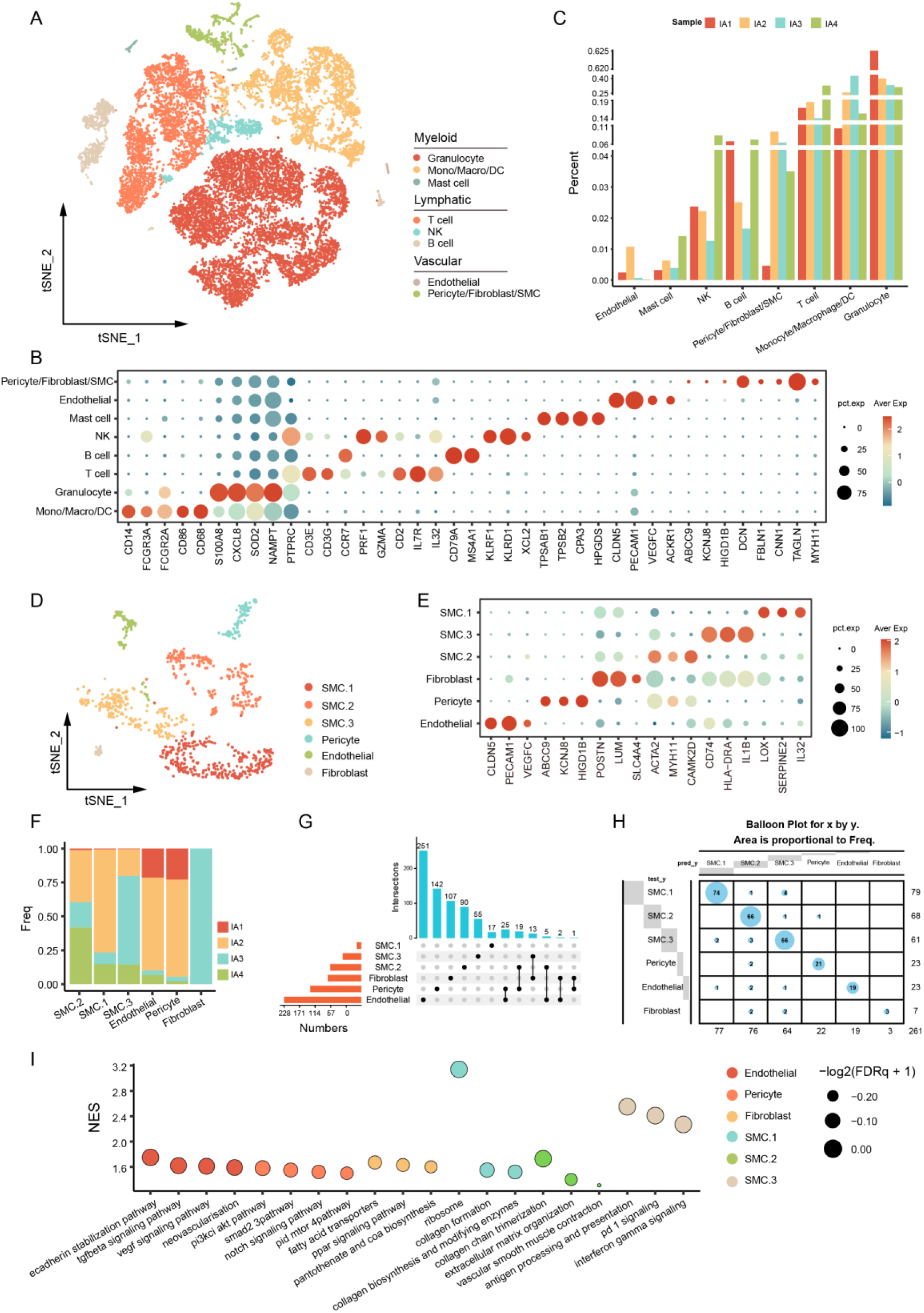
Cells and gene expression features of human IA tissue. (**A**) tSNE visualization of major cell types in IA wall. (**B**) Dot plot exhibiting conserved cell identity biomarkers. (**C**) Bar plot exhibiting the proportion of different cells in each sample. (**D**, **E**) Vascular cell types and states, and their unique gene markers. (**F**) Proportion of vascular cells in each sample. (**G**) The intersection of feature genes (log2FC > 1) of each cell type. (**H**) Random forest-based cell classifier for the identification of major cell types. (**I**) GSEA analysis of pathways enriched in each type/state of cell.

### Vascular cells in IA tissue

Normal cerebral arteries and arterioles are composed of endothelial cell, SMC, and perivascular fibroblast^8, 9^. Given that IA was characterized by pathological dilation of vascular wall and decellularization, the presence and status of these cells remain unclear^25^. To gain further insights, the vascular cells identified in the initial clustering were re-clustered at a higher resolution, with 4 types of vascular cells including endothelial cell (n = 89), pericyte (n = 92), fibroblast (n = 26), and smooth muscle cell (SMC, n = 826) being identified (Figure 1D).

Endothelial cells form the blood-facing surface of the intracranial artery, and is stressed and injured by the wall shear stress^26^. In this scenario, endothelial cells were marked by increased expression of CLDN5, CD31(PECAM1), and a recently identified endothelial marker gene, VEGFC^8^ (Figure 1E). Genes up-regulated in endothelial were mainly involved in the vascular development, tissue morphogenesis, cell migration and adhesion (Figure 1I, Supplementary Figure 2A).

Pericyte exists around microvessels and play a role in regulating blood flow and vessel formation and maintance^27, 28^. Although pericyte has been detected in the cerebral vasculature^8, 9^, we reported for the first time the presence of pericyte in human IA tissue. Pericyte was marked by ABCC9 and KCNJ8 and uniquely expressed HIGD1B (Pct.1 = 79.5%, Pct.2 = 1.9%, log_2_FC = 1.63), in line with previous study^8^. The identification of pericyte was based on two additional clues: 1. over 100 feature genes was found in pericyte which had little overlap with other cells (Figure 1G), and 2. the random forest-based cell classifier suggested that this type of cell was different from others (Figure 1H). Pericytes had activated signaling pathways including PI3K-Akt-mTOR, SMAD2, and NOTCH signaling (Figure 1E, Supplementary Figure 2), which are essential for cell survival and differentiation^29^. Previous study reported the presence of vasa vasorum in IA tissue^30^, cell-cell communication analysis found the interaction between endothelial and pericyte through PDGFB-PDGFRB and TGFB1-TGFBR2 pathway, and predicted the involvement of TGFB3 in the endothelial TGFB signaling activation (Supplementary Figure 2C-E), corroborating the role of pericyte in neovascularization^31^.

IAs are characterized by loss of SMCs^32^. We found that these cells can be further divided into three subgroups. SMC.1 up-regulated genes involved in extracellular matrix (ECM) portraying and inflammation (LOX, IL32, and SERPINE2), and had increased activity of signaling pathways associated with collagen biosynthesis and modifying (Figure 1E, I, Supplementary Figure 2F). SMC.2 highly expressed genes encoding actin and classical SMC transcription factor (ACTA2, MYH11, and CAMK2D), indicating the retention of contraction function. Meanwhile, SMC.2 participated in the ECM organization via the activation of signaling pathways associated with collagen production and organization (Figure 1E, I, Supplementary Figure 2G). SMC.3 was characterized by inflammation-associated genes (CD74, HLA-DRA, and IL1B), and the activation of signaling pathways including antigen presentation, interferon-gamma, and PD1 signaling, suggesting a pro-inflammatory role (Figure 1E, I, Supplementary Figure 2H). Similarly, three types of SMCs were identified by monocle, including SMC contraction (PDE3A), SMC synthetic 1 and 2 (TMSB10), and SMC inflammation (CD74, IL1B) (Supplementary Figure 2I).

Perivascular fibroblasts emerge early in the development of vessels, and plays a role in maintaining the vascular integrity^33^. The total number of fibroblasts captured and quantified was 26 and all were from the ruptured IA3 (Figure 1F), indicating that fibroblasts were scarce in IA tissue. Fibroblasts highly expressed POSTN, SLC4A4 and LUM and enriched in pathways involved in fatty acid metabolism (Figure 1E, I, Supplementary Figure 2), indicating that they were involved in ECM remodeling and inflammation.

### Immune cells in IA tissue

Awful amounts of immune cells were found in the IA tissue^32, 34^. These cells were re-clustered at a higher resolution, resulting in 2 major cell clusters and 18 subclusters (Figure 2A, B). Myeloid cells including neutrophils and monocyte/macrophages/DC made up a significant proportion (43.2% and 20.1%) and lymphoid cells including NK cell, T cell, B cell and subclusters to a less extend (31.2%) (Figure 2B, C). Each type of immune cell was detected in both ruptured and unruptured IA tissue (Figure 2A, C).

**Figure 2.**
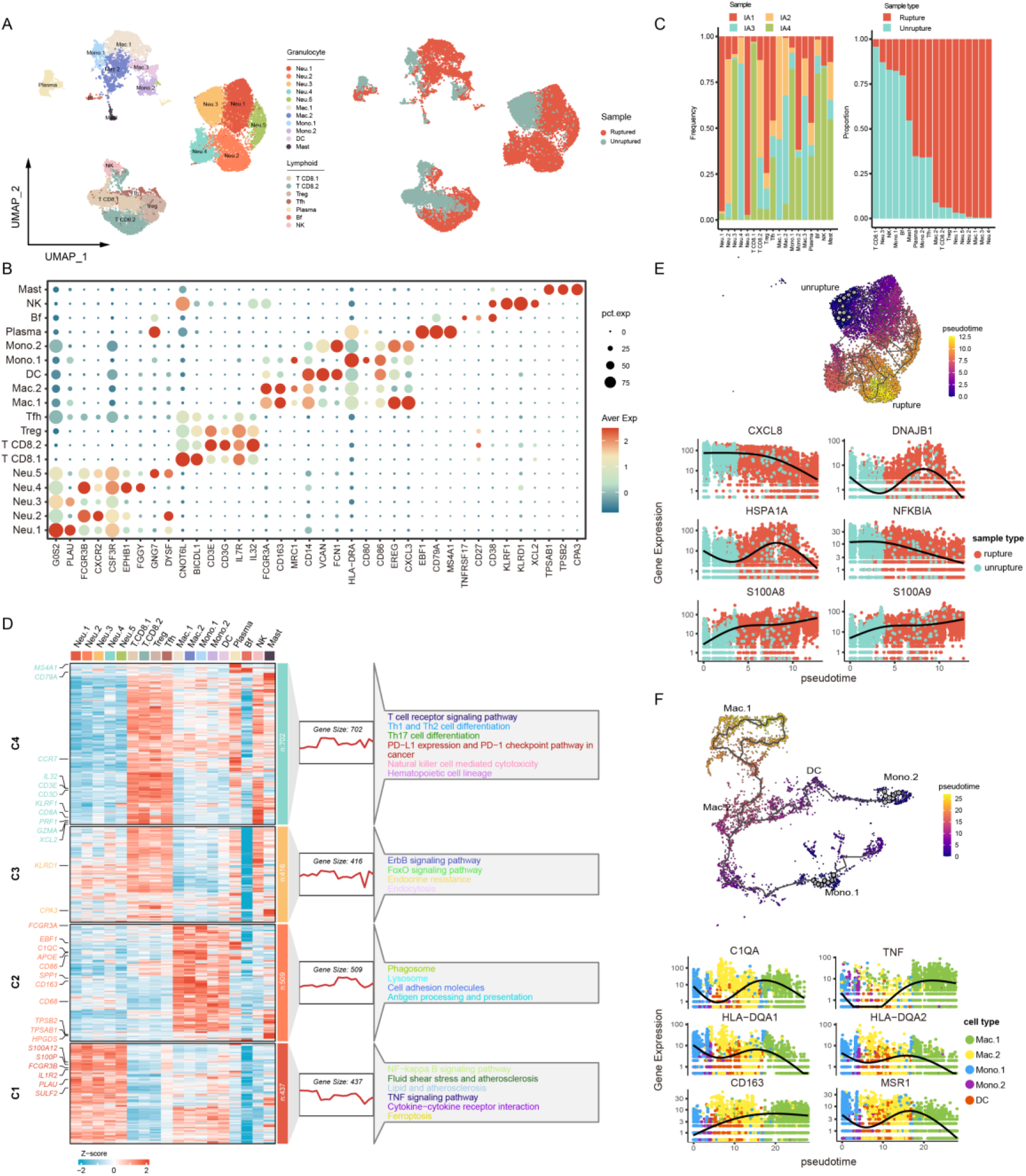
Immune cells and gene expression features of human IA tissue. (**A**) UMAP visualization of immune cells and their distribution in ruptured (red) and unruptured (green) IAs. (**B**) Dot plot showing conserved cell identity and state biomarkers. (**C**) Proportions of immune cells in each sample (left panel) and ruptured/unruptured IA tissue (right panel). (**D**) Gene expression patterns, feature genes, and enriched signaling pathways in the major immune cell types. (**E**) Alterations in neutrophil gene expression in ruptured and unruptured samples. The starting point of the trajectory was artificially set as cells in unruptured sample. (**F**) Alterations in gene expression in different types of monocyte/macrophages. The starting point of the trajectory was set as Mono.1 and Mono.2.

### Granulocytes in the IA tissue

Granulocyte, namely neutrophil (G0S2, PULA, FCGR3B) in this condition^35, 36^, was the most abundant immune cell in IA. The high proportion of neutrophils (31.1%) was found even in the unruptured sample. Genes highly expressed in neutrophils (Neu) were remarkably enriched in signaling pathways including NF-κB, TNF, and ferroptosis signaling (Figure 2D). Neutrophils are ferroptosis sensitive^37^ and expressed higher PTGS2 in contrast to other immune cells (Supplementary Figure 3A). Neutrophils were further divided into 5 subtypes (Figure 2A, B). Neu.1 was marked by increased expression of G0S2 and PLAU, and was active in chemotaxis, migration and inflammatory response (Figure 2B, Supplementary Figure 3B). Neu.2 highly expressed immunoglobulin receptor FCGR3B and CXCR2, and had enriched pathways associated with migrating and serine/threonine activity (Figure 2B, Supplementary Figure 3B). Neu.3 highly expressed eicosanoid metabolizing enzyme encoding gene ALOX5AP (log2FC = 1.65, Pct.1 = 0.94, Pct.2 = 0.53) and chemokine CXCL8 (log2FC = 1.55, Pct.1 = 1, Pct.2 = 0.73) (Figure 2B, Supplementary Figure 3C), indicating a role in synergistically biosynthesis of leukotrienes^38^. Neu.4 was characterized by the upregulation of EPHB1 that regulating cell migration and adhesion^39^, and FGGY that involved in energy metabolism (Figure 2B). Enrichment analysis also disclosed that Neu.4 was active in regulation of cell adhesion and fatty acid metabolic process (Supplementary Figure 3B). Neu.5 highly expressed G-protein beta-subunit GNG7 and DYSF that involved in membrane regeneration and repair, and genes involved in endoplasmic reticulum function and DNA damage repair like HSPH1 (log2FC = 2.96, Pct.1 = 0.98, Pct.2 = 0.50) and DNAJB1 (log2FC = 2.91, Pct.1 = 0.98, Pct.2 = 0.61), implying a subset of stressed neutrophil (Supplementary Figure 3B). The proportion of Neu.3 in unruptured IA tissue was particularly high (91.8%) (Figure 2C). When IA ruptured, neutrophils significantly up-regulated S100A8/9 that involved in the regulation of cell circle and inflammation, and down-regulated inflammation-related (CXCL8, NFKBIA) and genes involved in stress mitigation (HSPA1A, DNAJB1) (Figure 2E), suggesting a switch in neutrophil function when IA rupture.

Tissue infiltration of monocytes/macrophages occurs at an early stage of IA, and is of pathological significance^1, 40^. Monocytes/macrophages highly expressed CD14, FCGR3A, and type II MHC molecules and were defined as two types of monocytes (Mono.1/2), two types of macrophages (Macro.1/2), and DC. These cells were enriched in pathways including phagocytosis, lysosomal activation, and antigen presentation (Figure 2D). Mono.1 expressed both CD14 and FCGR3A, which may represent intermediate or non-classical monocytes, and had enriched pathways including antigen presentation, response to lipid, and positive regulation of immune response (Figure 2B, Supplementary Figure 3D). Mono.2 was CD14^+^, FCGR3A^-^, and had active inflammation response and cytokine production pathways (Figure 2B, Supplementary Figure 3D), thus may be classical monocytes. Mac.1 and Mac.2 highly CD163, with the former highly expressed CXCL3 and the latter CD206 (Figure 2B). Although both were involved in inflammation, Mac.1 was active in response to lipopolysaccharide and cytokine stimulus, Mac.2 was enriched in endocytosis and antigen presentation (Supplementary Figure 3D). DC up-regulated CD14, FCN1, and EGER, and had active dendrite morphogenesis pathway (Figure 2B, Supplementary Figure 3D). In the simulated evolutionary trajectory, there was significant upregulation of CD163, MSR1 and TNF, and downregulation of HLA molecules from monocytes to Mac.1 and Mac.2. Given the functional plasticity of macrophages, we further interrogated the functional status of Mac.1 and Mac.2, and defined macrophages with IFNg-induced polarization in vitro as classical activation and those with IL4-induced polarization as alternative activation. As a result, Mac.1 and Mac.2 both performed well in predicting classical and alternative activation expressed similar M1/M2 molecules (Supplementary Figure 3E, F). In addition, monocyte/macrophage/DC had generally increased ssGSEA score of bioactive lipids-related gene signatures (Supplementary Figure 3G), which was associated with alternative activation^14^. Together, these results indicated that the functional status of monocyte/macrophage/DC in IA was beyond M1 or M2 dichotomy.

A small amount of mast cells was identified, in line with the findings in mouse aneurysm model^10^. Mast cells in IA tissue highly expressed TPSAB1, TPSB2, and CPA3 that encoding tryptase and metalloprotease (Figure 2B). Enrichment analysis disclosed that mast cells promoted cell proliferation and activation and had high protein kinase and lipid metabolism activity (Supplementary Figure 3H). Besides, mast cells were involved in eicosanoid metabolism, and associated with platelet aggregation, indicating a pivotal role in local inflammation.

### Lymphocytes in the IA tissue

Lymphocytes are present in normal and aneurysmal vessel walls and involved in angiogenesis under certain circumstances^8, 9, 24, 41, 42^. Further, lymphocytes were re-clustered into 7 subclusters including T cells (CD8 T.1, CD8 T.2, T follicular helper (Tfh), and regulatory T (Treg)), B cells (plasma and B follicular (Bf)), and NK cell. CD8 T.1, NK, and Bf were abundant in unruptured IA, and once ruptured, the proportion of plasma cells, Tfh, Treg, and CD8 T.2 were elevated, indicating a dramatic perturbation in the properties of the inflammation.

CD8 T.1 and 2 had increased expression of CD3E, CD3G, and CD8A (log2FC = 1.06, Pct.1 = 0.52 in CD8 T.1, log2FC = 0.51, Pct.1 = 0.29 in CD8 T.2). CD8 T.1 was characterized by GZMK (logFC = 0.6) and granule secretory-related gene BICDL1 (Figure 2B). CD8 T.2 highly expressed cytotoxic markers including PRF1 (logFC = 0.56) and GNLY (logFC = 1.64). CD8 T.1 had activated lymphocyte activation and T cell receptor signaling pathway, and CD8 T.2 was enriched in T cell differentiation (Supplementary Figure 6A). During the simulated switching of status between the two cells, CD8 T.1 down-regulated S100A4 and up-regulated functional protein GZMB, and HSPA1B and DNAJB1 associated with stress or senescence (Supplementary Figure 4B). Treg was characterized by the expression of IL2RA (log2FC = 0.50) and SELL (log2FC = 0.44) and were involved in inducing cell apoptosis and humoral immune response (Supplementary Figure 4A). Tfh was characterized by ICOS (log2FC = 0.41) and IL21R (log2FC = 0.49). Interestingly, Tfh not only participated in T cell differentiation, but also regulating muscle cell differentiation (Supplementary Figure 4A). Cellchat predicted two patterns of interaction between Tfh and SMCs (Supplementary Figure 4C, D), in which secreted nicotinamide phosphoribosyl transferase (NAMPT) plays a role in the survival of SMC^43, 44^.

B cells were defined as plasma cell and Bf. Plasma cell was characterized by CD79A and MS4A1 and had activated B cell receptor pathways. Plasma cells were involved in antigen presentation and T cell differentiation (Figure 2B, Supplementary Figure 4E). Bf highly expressed TNFRSF17 and CD38 (log2FC = 0.86, Pct.1 = 0.50) and was involved in immune response, antigen presentation, and angiogenesis (Supplementary Figure 4F). A small amount of NK cells was identified and highly expressed KLRF1, KLRD1, and XCL2 (Figure 2B). NK cells up-regulated GZMB (log2FC = 1.98, Pct.1 = 0.91, Pct.2 = 0.04) and PRF1 (log2FC = 1.48, Pct.1 = 0.85, Pct.2 = 0.06), which may aggravate IA through a direct cytotoxicity.

### Inflammatory features associated with IA rupture

To further explore inflammation features associated with IA rupture, two external microarray-based datasets were collected. The ssGSEA scores of Mac.1, Mac.2, DC, and Mono.2 were robustly increased in the ruptured IA tissue, indicating an increased cell abundance (Figure 3A, B). ROC analysis disclosed that the ssGSEA score of Mac.1 gave the best performance in predicting IA rupture (Figure 3C, D), indicating an association between Mac.1 and IA rupture. These results were in accordance with previous findings that specific type of monocyte/macrophage was implicated in bleeding of cerebral arteriovenous malformations and thoracic aortic aneurysm^8, 11^. Given the phenotypic sophistication of Mac.1, further analysis was conducted to deconstruct transcriptional network of these cells. Several transcriptional factors and corresponding high-confidence target genes were identified, and Mono.1 and Mac.1 shared similar transcriptional regulatory patterns (Figure 3E, Supplementary Figure 5A). NF-κB (NFKB1, NFKB2 and RELB) was activated in Mac.1, together with its negative regulator HIVEP2 (Figure 3E). Accordingly, the transcriptional networks of Mac.1 and Mac.2 were distinct, with the high-confidence up-regulated target genes in Mac.1 enriched in the TNF signaling, contrary to that of Mac.2 (Supplementary Figure 5B-D). These results suggested that Mac.1 was pro-inflammatory orientated and Mac.2 the opposite. Moreover, WGCNA algorithm identified the intrinsic gene expression pattern of Mac.1, some of which were involved in aneurysm progression and rupture (Supplementary Figure 6), corroborating its role in IA rupture.

**Figure 3.**
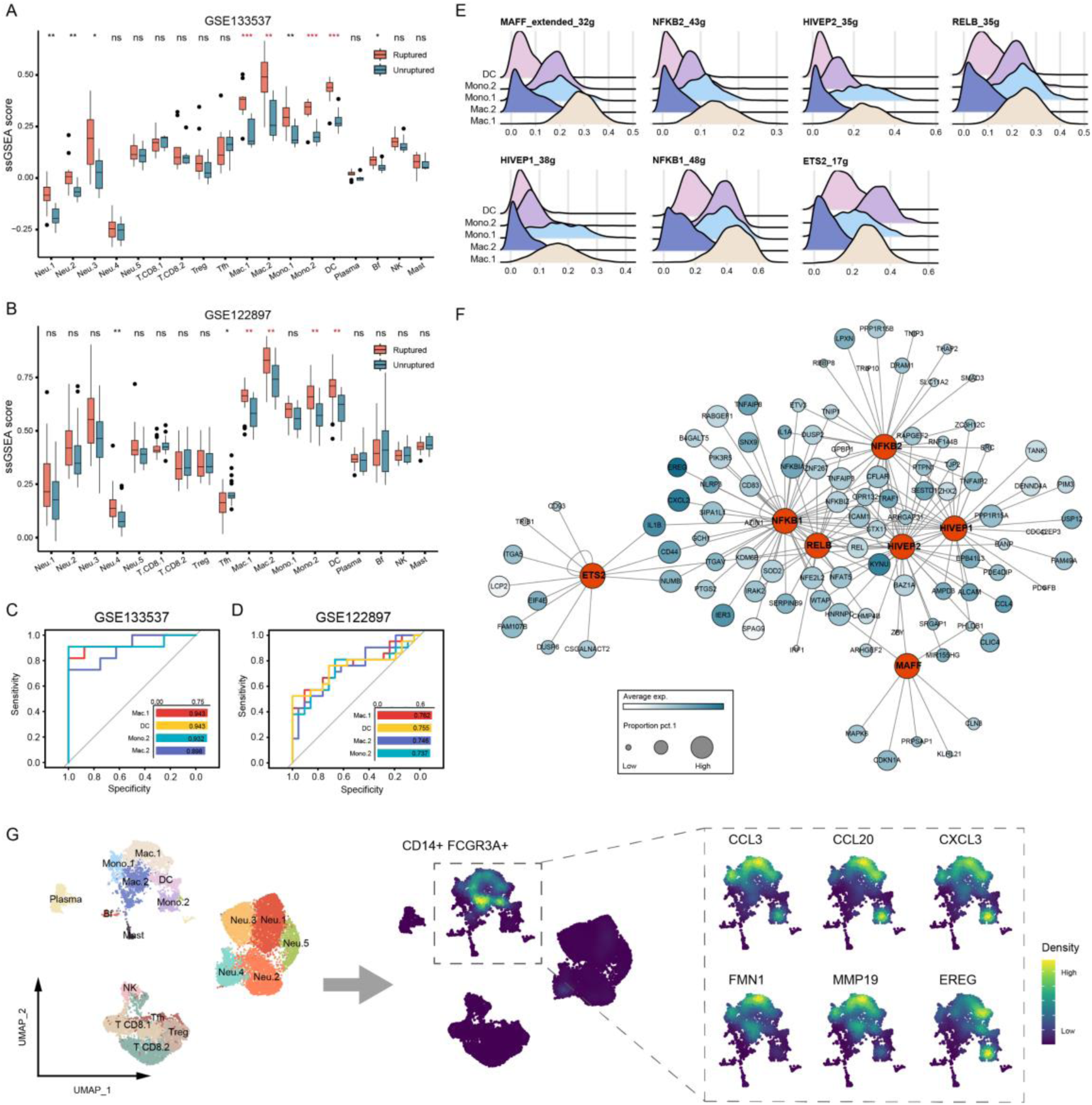
Identification of rupture-associated macrophage and its transcriptomic features. (**A, B**) ssGSEA score of immune cells in ruptured and unruptured IA tissues. Genes with log2FC > 0.5 and Pct.2 < 0.15 in each cell type were used as signature genes. (**C**, **D**) Performance of gene signatures in predicting IA rupture. (**E**) The activation pattern of transcriptional factors mainly associated with Mac.1 and Mono.1. (**F**) The network of transcriptional factors mainly activated in Mac.1 and their high-confidence target genes. (**G**) Density plot showing feature genes mainly expressed by Mac.1.

To interrogate how Mac.1 may facilitate IA rupture, the interactome between Mac.1 and vascular cells were estimated by cellphoneDB. As a result, the regulation of vascular cells (including endothelial, pericyte, SMC, and fibroblast) by Mac.1 was largely dependent on the secretion of TNFα (encoded by TNF) (Figure 4A, B). TNFα is functionally pleiotropic depending on the receptor it binds, for instance, binding to TNFR2 (encoded by TNFRSF1B) is involved in cell survival, while binding to TNFR1 (encoded by TNFRSF1A) or FAS may induce apoptosis^45^. The receptors associated cell death or survival showed a cell type heterogeneity (Figure 4C). For example, FAS was mainly expressed by SMC.2, NK and some CD8 T cells, and TNFRSF1A was expressed by most vascular cells (SMC.2, pericyte, endothelial, and fibroblast) and immune cells like neutrophil. In turn, the expression of TNFRSF1B was higher than TNFRSF1A in immune cells, indicating that TNFα mainly maintains the survival of immune cells and induces death of vascular cells. On the other hand, Mac.1 interacted with other immune cells through the binding of different ligand-receptor pairs (Figure 5A, B). In terms of significant ligand-receptor interactions, Mac.1 had the highest counts of both ligands and receptors (Figure 5C), indicating that Mac.1 should be a hub node in the chronic inflammation of IA tissue.

**Figure 4.**
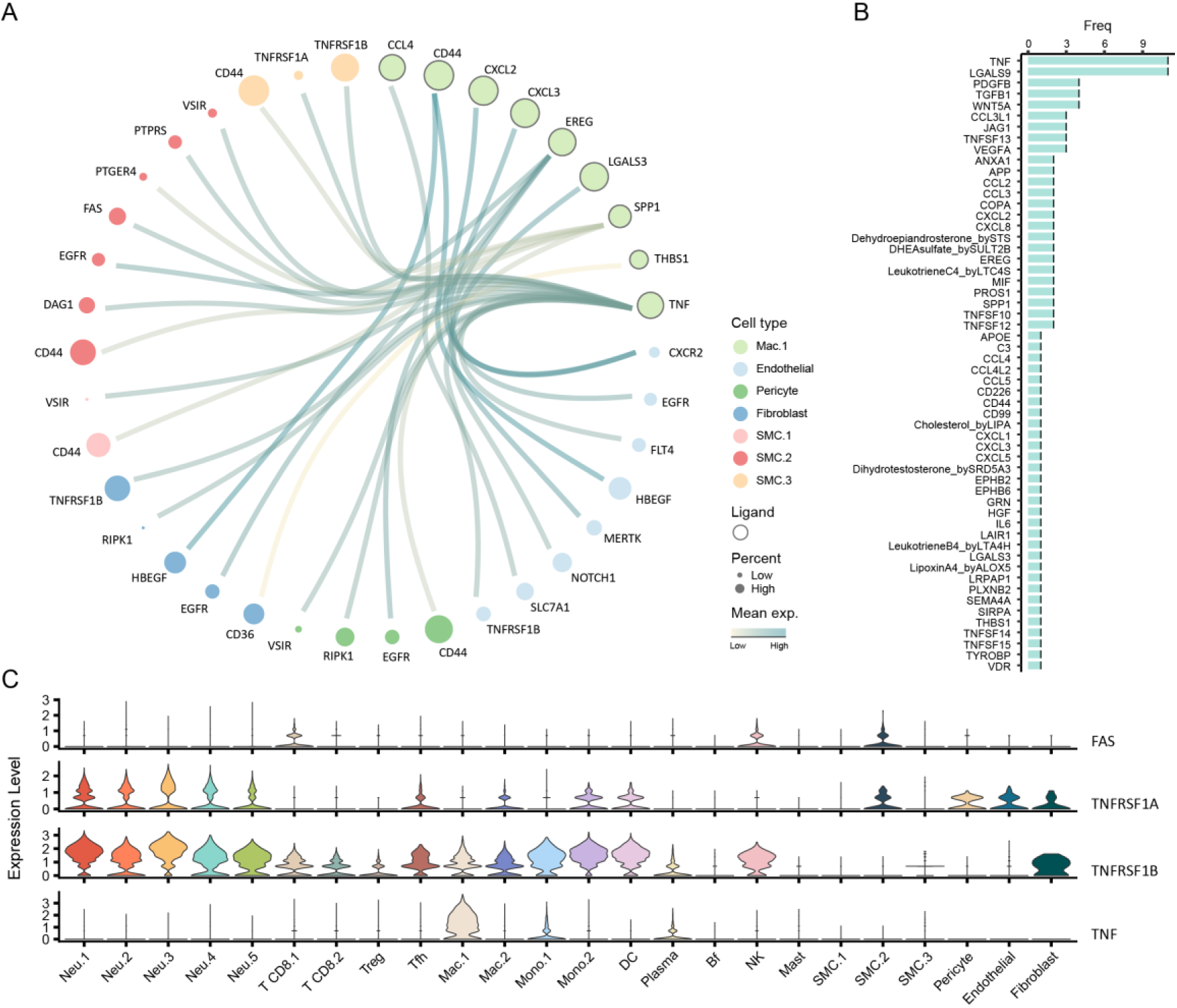
Interactions between rupture-associated macrophage and vascular cells. (**A**) Circus plot exhibiting top expressed ligands of Mac.1 and their binding receptors expressed by vascular cells. Bubble size is proportional to the percentage of cells expressing corresponding gene. The color of line was associated with the mean expression of ligand-receptor pairs estimated by cellphoneDB. (**B**) Frequency of ligands of Mac.1 that interact with vascular cells. (**C**) Expression of ligands and receptors associated with TNFα-mediated cell survival and death.

**Figure 5.**
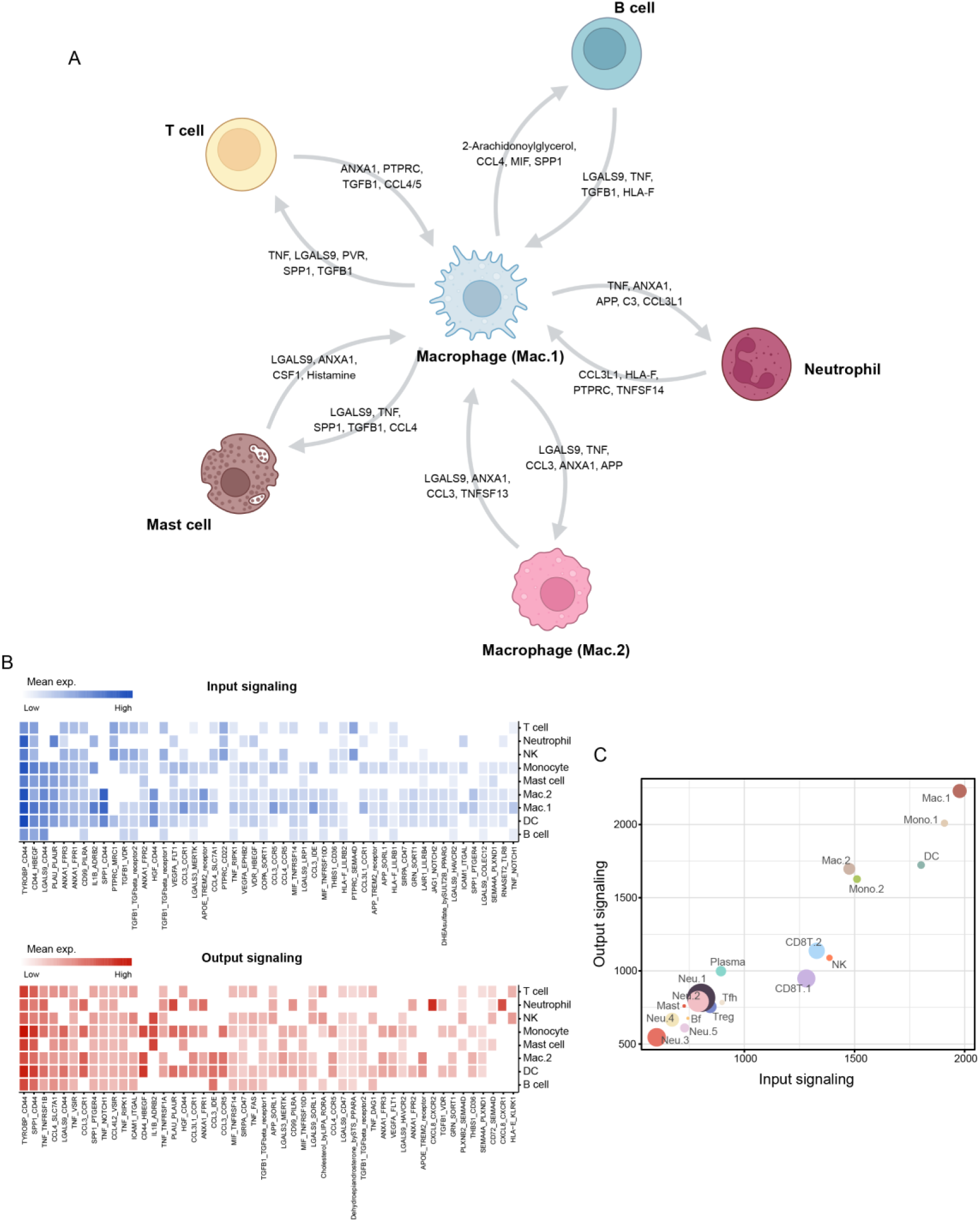
Interactions between rupture-associated macrophage and other immune cells. (**A**) Top ligands mediating immune cells interactions. (**B**) Top 50 ligand-receptor pairs mediating interactions between Mac.1 and other immune cells. The input signaling means the binding of ligands expressed by other cells to Mac.1 receptors, the output signaling is the opposite. (**C**) Cell communication strength. Statistically significant input and output signal counts of each type of immune cell estimated by cellphoneDB. The x-axis is the count of receptors (input) and the y-axis the ligands (output). Size of the bubble is proportional to the proportion of each type of cell.

## Discussion

Intracranial aneurysm progression is driven by multifaceted contributions. This study provided a detailed cell atlas of human IA tissue and highlights the pathological implication of inflammation. We expanded cellular diversity of human IA by the identification of pericyte in addition to the previously reported vascular cells, and provided a compendium of immune cell types and corresponding functional characteristics. Notably, we identified a subset of macrophage that was closely associated with IA rupture, with inflammatory activity being its major functional orientation. The CCL3/CXCL3 macrophage acted as a hub of local inflammatory network and may induce vascular cell death in a TNFα-dependent manner.

Recent studies thoroughly characterized the major cell types and corresponding transcriptome features of the cerebrovascular system in near-physiological states, including endothelial, smooth muscle cell, pericyte, perivascular fibroblast, and less common fibromyoblast^8, 9^. Endothelial can be further divided into arterial, venous and capillary subtypes with distinct molecular markers. These major cell types also exist in human IA tissue. Interestingly, we for the first time identified pericyte in IA tissue, which according to previous studies exists in small vessels or capillaries^27, 46^. Pericyte is morphologically divided into reticular and fascicular subtypes, and play a role in regulating blood flow and maintaining the brain-blood barrier^46^. Endothelin and PDGFBR expressed by endothelial recruit bone marrow-derived pericytes, which, in turn, promote endothelial cell differentiation and maturation via the TGFβ pathway and participate in neovascular sprouting^31, 47^. Recently, vasa vasorum has been identified in human IA tissue, which coincides with the scenario of endothelial-pericyte interaction-mediated neoangiogenesis^30, 31, 47, 48^. Nevertheless, lymphatic vessel, another microtubular system, also exists in the wall of saccular IA^49^. The lymphatic vessels were characterized by LYVE-1 and podoplanin positivity staining and the presence of SMC peripherally, with the latter expressing both αSMA and EGFR3. These results mean that the small duct-like structures found in the IA wall need to be carefully identified to further explore the function and potential pathological significance of pericyte.

Astounding number of immune cells were identified in both unruptured and ruptured IA tissues, consistent with previous histology-based findings^4, 5, 50^. Immune cells including T and B lymphocytes, monocyte/macrophage, neutrophil, and to a lesser extent NK cell, mast cell, etc. are involved in IA rupture^10, 41, 50, 51^. Particularly, monocyte/macrophages are widely present in IA tissues and can be found at an early stage of the disease^4, 10, 41^. The level of monocyte/macrophages positively correlates with the inflammatory response in the IA wall and surrounding tissues, and increases the risk of SAH^52, 53^. Besides, monocyte/macrophages have also been identified as responsible for the hemorrhage of intracranial arteriovenous malformation and thoracic aortic aneurysm (TAA)^8, 11^, making these cells a potential therapeutic target for unruptured IA. Previous studies, and our results, have shown that the transcriptome heterogeneity of macrophages^14^. Thus, the question at hand is which type of macrophage should be actually targeted. Active inflammatory response activity is a distinctive feature of macrophages leading to AVM and TAA rupture, which is closely associated with transcriptional activation of classical and non-classical NF-κB signaling pathway^54–56^. Indeed, we found that CCL3/CXCL3 macrophages associated with IA rupture were manifested as transcriptional activation of NF-κB. Besides, these cells had an activation of HIVEP2, which acts as a braker of the immune response through inhibiting NF-κB^57, 58^. Given the potential role in induing vascular cell death and orchestrating inflammation, CCL3/CXCL3 macrophages were a promising therapeutic target for unruptured IA, where HIVEP2 may be a plausible molecule to manipulate their inflammatory activity.

Morphologically, saccular and fusiform aneurysms are common while serpentine and blood blister-like aneurysms (BBA) are less common. Different appearances of IA may be associated with different etiological underpinnings, but factors predisposing to IA progression can be summarized in two categories: direct damage to the IA wall from high wall shear stress and low wall shear stress accompanying inflammation-mediated IA wall remodeling^1, 59^. These features can be glimpsed in histological studies, with the limitations of the size of usual IA. Theoretically, the progression of IA due to high wall shear stress and inflammation should have distinct histological features, at least in the proportion and type of immune cells in the IA dome. Current knowledge of the characteristics of small size saccular IAs that may be in an early pathological stage is inadequate due to the difficulty of sampling^5^. In fact, in the initial phase of the study we found that it is difficult to obtain sufficient numbers of cells from these tissues, probably due to the sparseness of vascular cells and the low level of immune cell infiltration. Thus, samples withstand quality inspection were those with larger and thicker-walled IAs, which were more like to be affected by inflammation. Similarly, despite the extremely thin wall of BBA, Wen et al. still identified a large number of immune cells, corroborating that the inflammation was the underlying factor in the rupture of some IAs encountering less hemodynamic perturbation.

In summary, we compiled a catalogue of cells and their gene expression features of IA wall through single-cell RNAseq of general human IA samples, and identified a type of macrophage as promising therapeutic for unruptured IA. Due to the importance of inflammation and current paucity in understanding of the impact of immune cells on IA, more studies could be devoted to revealing how other immune cells affect the integrity of the IA wall.

## Author contribution

Study design: Yi Liu, Hang Ji, and Yue Li. Data collection and curation: Yue Li, Haogeng Sun, Ruiqi Chen, and Anqi Xiao. Data analysis: Hang Ji, Ran Zhou, and Yue Li. Manuscript drafting: Hang Ji and Yue Li. Manuscript revise: Rong Wang, Yongbo Yang, and Chao You. Study supervising: Yi Liu and Chao You. Hang Ji and Yue Li contributed equally to this study.

## Data availability statement

Data can be obtained from the corresponding author upon reasonable request.

## Supplementary Tables

**Supplementary Table 1. Differentially expressed genes of immune cells.**

**Supplementary Table 2. Positive and negative hits of IL-4- and IFNγ-polarized macrophages.** Positive hits are intersections of upregulated genes in macrophages after 6, 12, 24 and 72 hours of incubation with the corresponding conditions, negative hits are the intersections of downregulated genes.

**Supplementary Table 3. Gene signatures of different conditions-polarized macrophages.**

## Supplementary Figure Legends

**Supplementary Figure 1.**
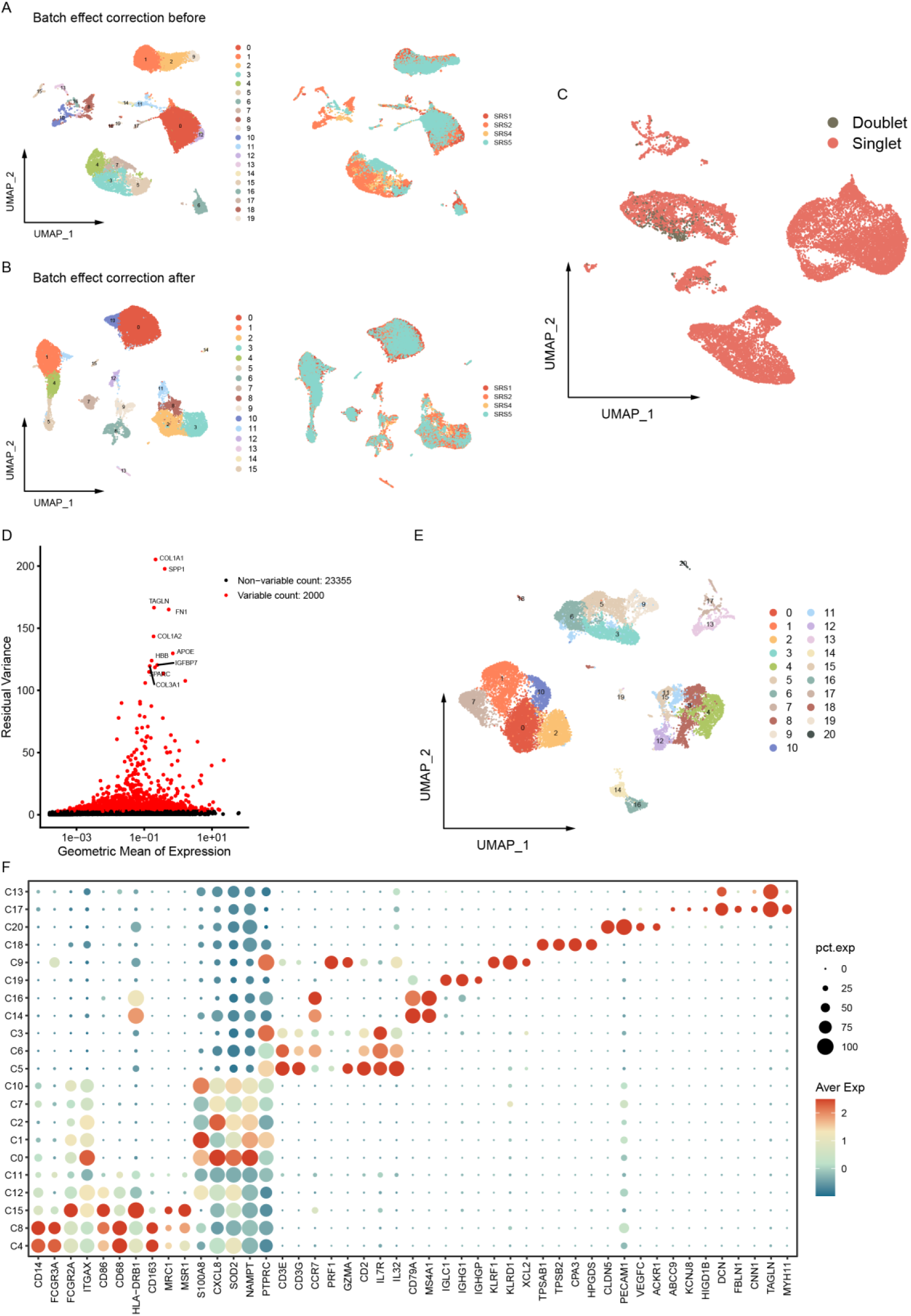
Data pre-processing and preliminary clustering. (**A**, **B**) Before and after batch effect correction. (**C**) Identification of doublets. (**D**) Estimation of highly variable genes. (**E**, **F**) Preliminary cell clustering and marker genes of each cluster.

**Supplementary Figure 2.**
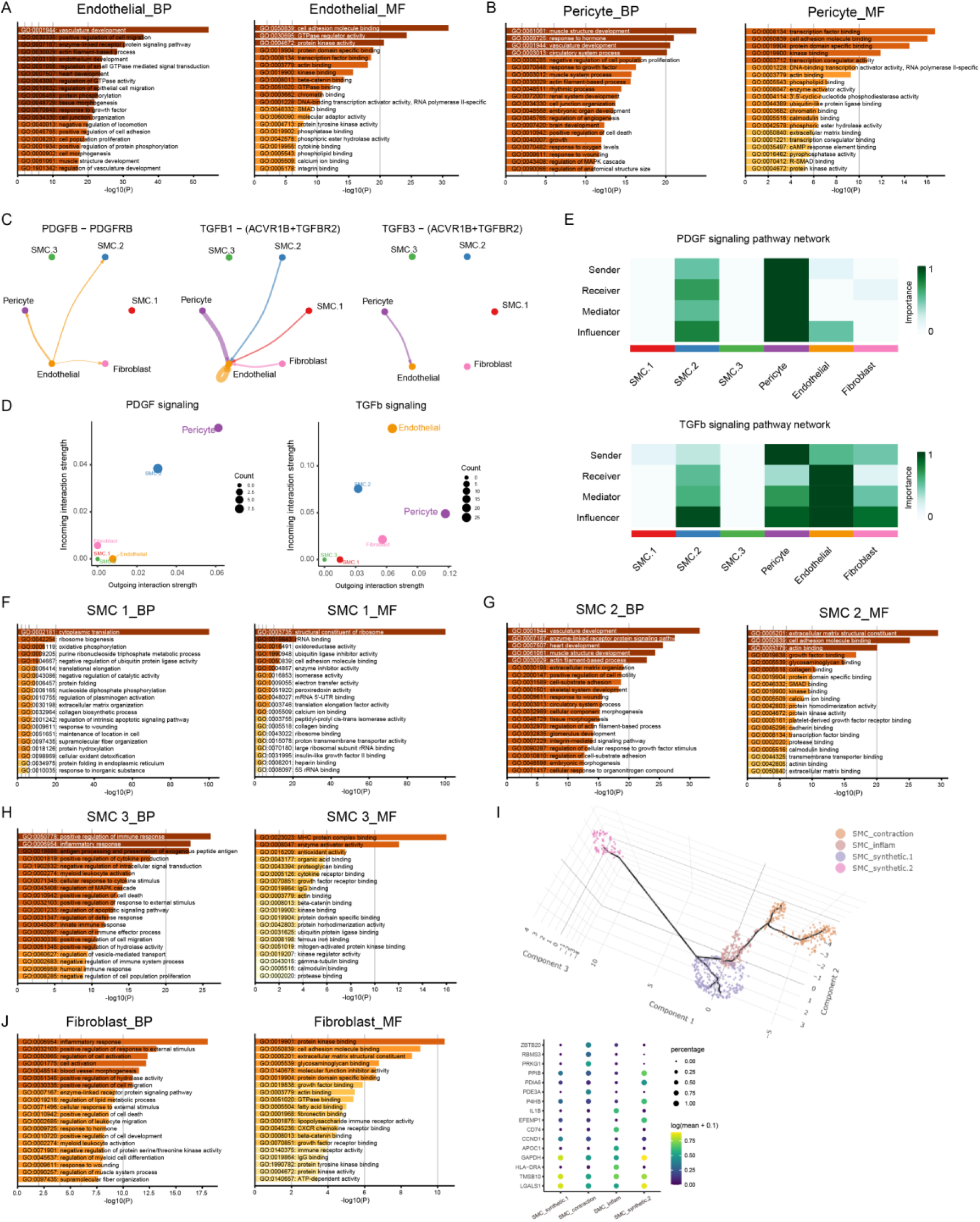
Gene expression features of vascular cells. (**A**, **B**) Gene ontology (GO) enrichment analysis of feature genes of endothelial and pericyte. The cut-off of feature genes was set as log2FC > 0.85 and Pct.1 > 0.75. (**C**) Cell-cell communication based on the Cellchat algorithm. The interactions between PDGFB, TGFB1, and TGFB3 with their receptors were exhibited. (**D**) Dot plot exhibiting the strength of PDGF and TGF-β signaling pathways in each type of vascular cell. (**E**) The role of vascular cells in the PDGF and TGF-β signaling pathways. (**F**, **G**, **H**) GO enrichment analysis of feature genes of SMCs. The cut-off of feature genes was set as log2FC > 0.85 and Pct.1 > 0.75. (**I**) The subtypes of SMC identified using the monocle algorithm and their feature genes. Inflam: inflammation. (**J**) GO enrichment analysis of feature genes of fibroblast. Cut-off of feature genes were the same to above.

**Supplementary Figure 3.**
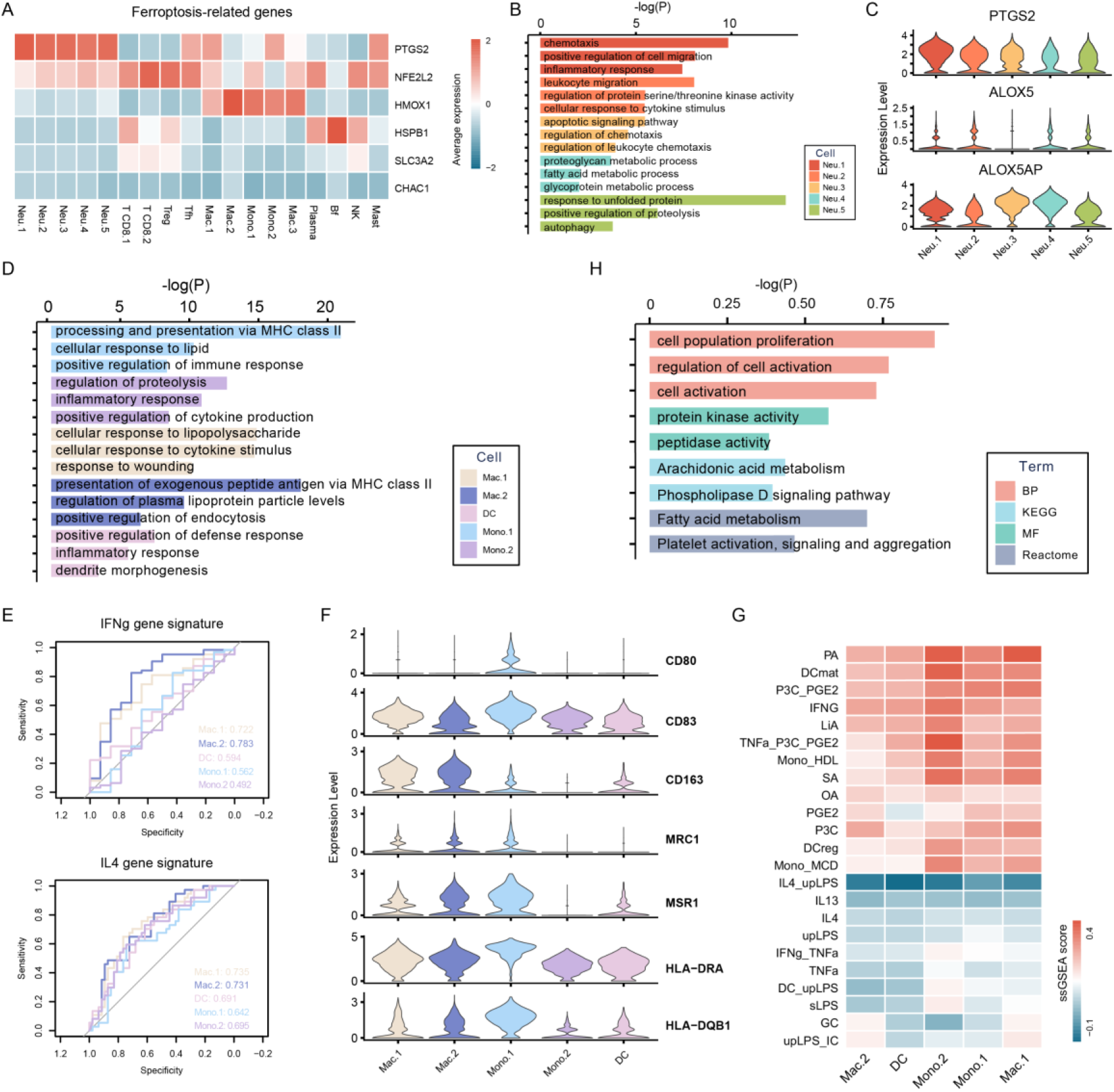
Gene expression features of myeloid cells. (**A**) Heatmap showing the expression of ferroptosis-related genes of each type of immune cell. (**B**) GO and Kyoto Encyclopedia of Genes and Genomes (KEGG) enrichment analysis of feature genes. Only top 3 terms were exhibited. The cut-off of feature genes was set as log2FC > 0.85 and Pct.1 > 0.75. (**C**) The expression of genes encoding cyclooxygenase and lipoxygenase in neutrophils. (**D**) GO and KEGG analysis of feature genes of monocyte/macrophage/DC. (**E**) The performance of feature genes of each type of mono/macro in predicting the IFN-γ and IL-4 gene signatures. Feature genes were defined as those with log2FC > 0.85 and Pct.1 > 0.75. IFN-γ and IL-4 gene signatures are conserved genes of macrophage induced by corresponding conditions in vitro at different timepoints (Supplementary Table 2). Upregulated genes were set as positive hits and downregulated negative hits. (**F**) The expression of conventional macrophage markers. (**G**) Heatmap showing the ssGSEA score of different stimuli-induced macrophage gene signatures in each type of mono/macro (Supplementary Table 3).

**Supplementary Figure 4.**
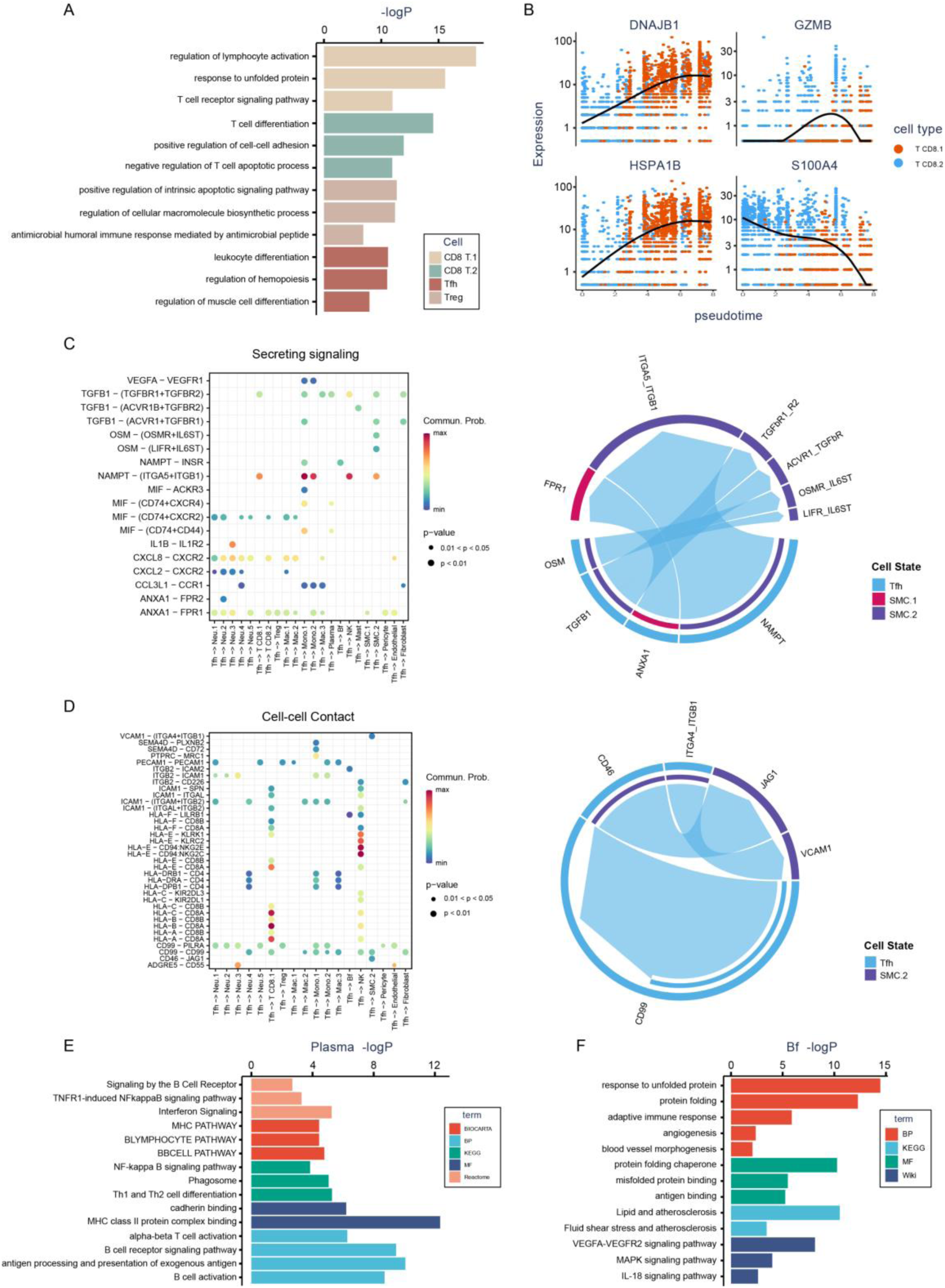
Gene expression features of lymphocytes. (**A**) GO and KEGG analysis of T cells. (**B**) Gene expression alterations between the two subtypes of CD8 T cells. (**C**, **D**) The output signaling from Tfh to other cells through secreting and directly contact manner. (**E**, **F**) Enrichment analysis of feature genes of plasma and follicular B cell. The cut-off of feature genes was set as log2FC > 0.85 and Pct.1 > 0.75.

**Supplementary Figure 5.**
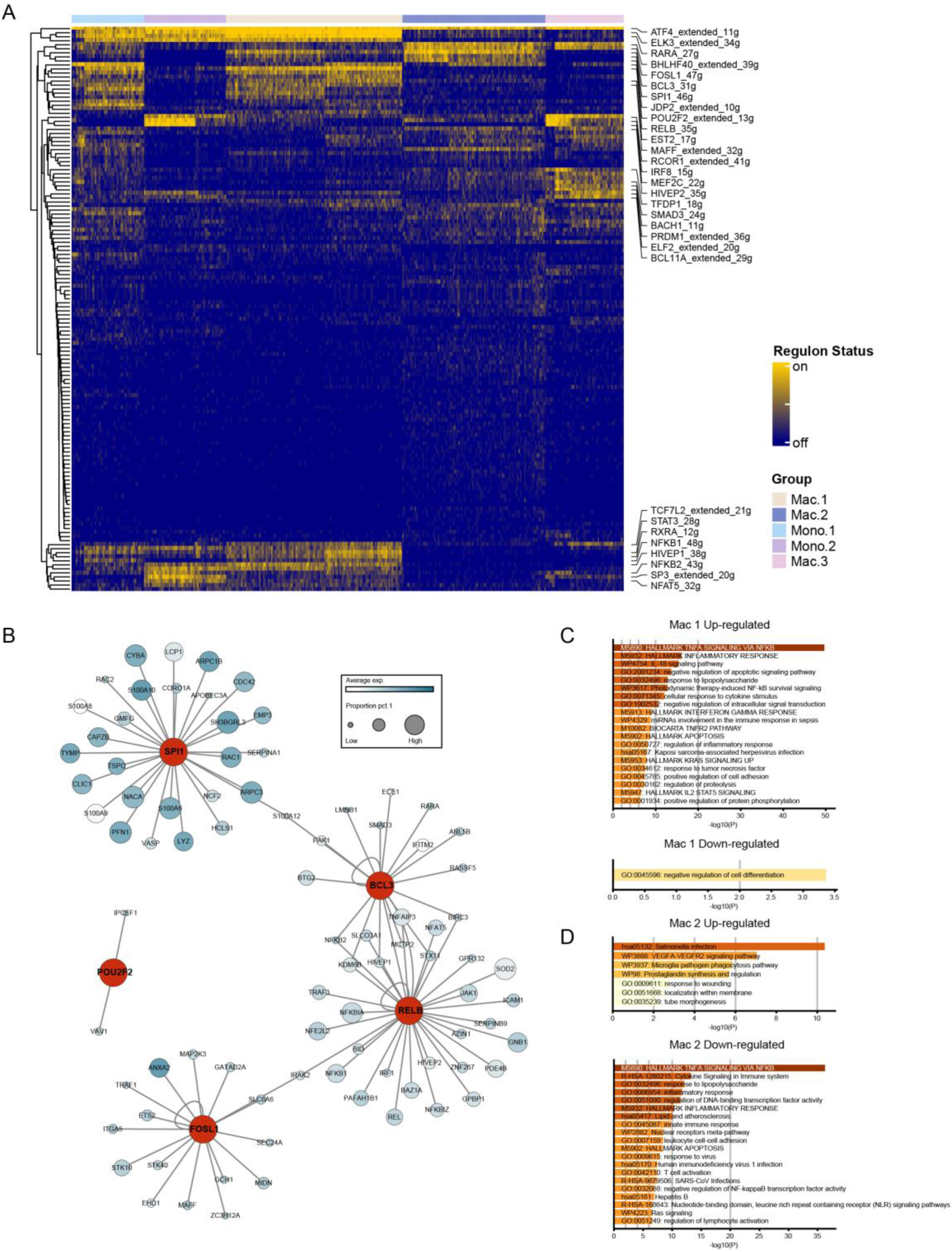
Transcriptional features of monocyte/macrophages. (**A**) Heatmap showing the activity of transcription factors in each type of cell, labelled by the algorithm as on (activated) and off (in-activated) state. Transcriptional factors differentially activated between cells were labelled out. (**B**) The activated transcriptional factors and corresponding high-confidence target genes of Mac.2. (**C**, **D**) GO analysis of high-confidence target genes of Mac.1 and Mac.2.

**Supplementary Figure 6.**
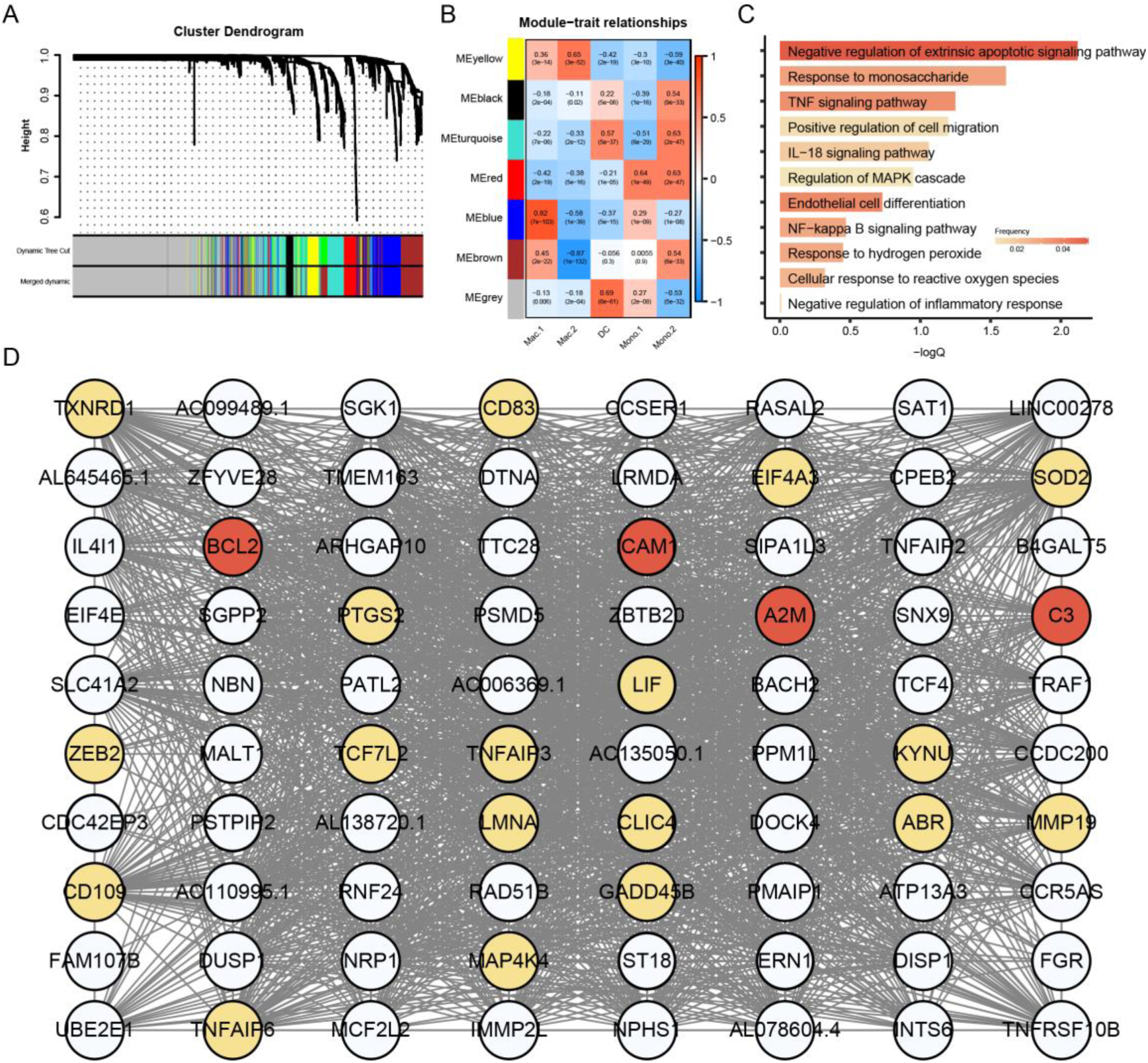
The intrinsic gene expression pattern of Mac.1. (**A**) Dendro plot of clusters of pseudo-cells. (**B**) Gene modules and their correlation with cell types. (**C**) GO and KEGG analysis of genes in module blue. (**D**) Network of hub genes of the module blue. Hub genes were selected using Kwithin.

